# Neural responses to broadband visual flicker in marmoset primary visual cortex

**DOI:** 10.1101/2022.02.05.479227

**Authors:** Jakob C. B. Schwenk, Maureen A. Hagan, Shaun L. Cloherty, Elizabeth Zavitz, Adam P. Morris, Nicholas S. C. Price, Marcello G. P. Rosa, Frank Bremmer

## Abstract

Temporal information is ubiquitous in natural vision and must be represented accurately in the brain to allow us to interact with a constantly changing world. Recent studies have employed a random stimulation paradigm to map the temporal response function (TRF) to luminance changes in the human EEG. This approach has revealed that the visual system, when presented with broadband visual input, actively selects distinct temporal frequencies, and retains their phase-information for prolonged periods of time. This non-linear response likely originates in primary visual cortex (V1), yet, so far it has not been investigated on a neural level. Here, we characterize the steady-state response to random broadband visual flicker in marmoset V1. In two experiments, we recorded from i) marmosets passively stimulated under general anesthesia, and ii) awake marmosets, under free viewing conditions. Our results show that LFP coupling to the stimulus was broadband and unselective under anesthesia, whereas in awake animals, it was restricted to two distinct frequency components, in the alpha and beta range. Within these frequency bands, coupling adhered to the receptive field (RF) boundaries of the local populations. The responses outside the RF did not provide evidence for a propagation of stimulus information across the cortex, contrary to results in human EEG studies. This result may be explained by short fixation durations, warranting further investigation. In summary, our findings show that during awake behavior V1 neural responses to broadband information are selective for distinct frequency bands, and that this selectivity is likely controlled actively by top-down mechanisms.

## Introduction

The accurate representation of temporal information is a critical part of all sensory processing. In the visual domain, this information can be conveyed in feature dimensions, the most reliable of which to evoke neural responses experimentally is luminance contrast. Luminance-modulated images have been utilized in EEG research for decades by the use of steady-state visual evoked potentials (SSVEP), i.e. the periodic responses to prolonged visual flicker at a constant temporal frequency (e.g., Herrmann 2001; for a comprehensive review, see Vialatte et al. 2010). Among other applications, SSVEPs have become a major tool in the investigation of visual attention (Morgan et al., 1996) and working memory (Perlstein et al., 2003; Ellis et al., 2006), and have been employed as a signal to control Brain-Computer-Interfaces (Fazel-Rezai et al., 2013; Abiri et al., 2019). Similar stimulation protocols have been used to characterize the temporal response properties of V1 neural populations in animal models (Rager and Singer, 1998). The visual stimuli used in the above studies, however, always contained only a single constant frequency. While this allows for tight control over the desired response, it does not reflect the temporal information that is available in natural vision, which, in most contexts, is not fully predictable and comprises a broader spectrum of temporal frequencies. VanRullen and MacDonald (2012) addressed this limitation by introducing a variation of the visual flicker paradigm which uses broadband random luminance sequences. By reverse-correlation of recorded EEG with the presented sequences, they showed that the visual impulse-response in the human EEG contains two components: a transient, relatively broadband response within the first 150-200 ms, followed by sustained (up to > 1 sec lag) coupling at the individual alpha frequency. This narrow-band component (the “perceptual echo” response) shows a spatio-temporal distribution that strongly suggests propagation across the cortex in the form of a traveling wave (Lozano-Soldevilla and VanRullen, 2019). Recent studies have built upon this finding by using the same paradigm to identify possible functional roles of the echo component in active rhythmic sampling and prediction of visual input (Brüers and Vanrullen, 2017; Chang et al., 2017; Gulbinaite et al., 2017; Benedetto et al., 2018; Alamia and VanRullen, 2019; Schwenk et al., 2020).

Despite this growing interest, the neural basis of the underlying response(s) remains unknown, with the human EEG studies presenting only sparse insight in this direction. The initial findings by VanRullen and MacDonald (2012) indicate that the full response is likely based on a non-linear interaction of the temporal response properties of local V1 populations (in the sense that it cannot be modeled as the sum of responses to stimulation at single frequencies). In the steady state that is reached within a few hundred ms after onset of the stimulus-sequence, the response seems to be driven mainly by phase-coupling of the EEG to the stimulus (Chang et al., 2017; Schwenk et al., 2020) in the same way as the SSVEP (Moratti et al., 2007). This may point towards a sparse temporal coding regimen in which the activity of V1 cells is adapted and temporal information is relayed mainly by spike timing. Lastly, the evidence for a wave-like propagation of the echo component implies that the response extends to regions outside the retinotopic representation of the stimulus. This poses the question of whether the alpha-band selection and separation into response components occurs already at the retinotopic site, or requires signal summation over larger volumes of cortex.

The above considerations clearly illustrate that a characterization of neural responses to broadband luminance flicker using intracortical recordings is needed. Towards this aim we recorded with electrode arrays from the primary visual cortex (area V1) of marmoset monkeys, using the stimulation procedure introduced by VanRullen and MacDonald (2012). To establish whether the selection of stimulus information depends on active visual behavior, we compare data obtained in experiments from anesthetized (exp. 1) and awake, behaving animals (exp. 2).

## Methods

All experimental procedures were approved by the Monash Animal Research Platform Animal Ethics Committee and carried out in accordance with the Australian Code of Practice for the Care and Use of Animals for Scientific Purposes.

### Experiment 1: Surgical and experimental procedures

Recordings were performed during acute experiments in two marmoset monkeys (*Callithrix jacchus*, male/female, cj12 and cj11). The preparation for electrophysiological studies of marmosets has been described previously (Yu and Rosa, 2010). Briefly, injections of atropine (0.2 mg/kg, i.m.) and diazepam (2 mg/kg, i.m.) were administered as premedication, 30 minutes prior to the induction of anesthesia with alfaxalone (Alfaxan, 10 mg/kg, i.m., Jurox, Rutherford, Australia), allowing a tracheotomy, vein cannulation and craniotomy to be performed. The animal received an intravenous infusion of pancuronium bromide (0.1 mg/kg/h; Organon, Sydney, Australia) combined with sufentanil (6 μg/kg/h; Janssen-Cilag, Sydney, Australia) and dexamethasone (0.4 mg / kg / h; David Bull, Melbourne, Australia), and was artificially ventilated with a gaseous mixture of nitrous oxide and oxygen (7:3). This regime ensures long-term anesthesia with less suppression of early response components in primary sensory areas, in comparison with isoflurane or barbiturates (Rajan et al., 2013). The electrocardiogram and level of cortical spontaneous activity were continuously monitored. Administration of atropine (1%) and phenylephrine hydrochloride (10%) eye drops (Sigma Pharmaceuticals, Melbourne, Australia) resulted in mydriasis and cycloplegia. Appropriate focus and protection of the corneas from desiccation were achieved by means of hard contact lenses selected by streak retinoscopy.

A craniotomy was performed to allow access to the left occipital cortex for the implantation of a 10 × 10, 96-channel Utah array (1.5 mm in length, spaced at 400 μm intervals, with platinum electrode sites, Blackrock Microsystems, Salt Lake City, USA). Extracellular neural data were recorded using a Cerebus System (Blackrock Microsystems, Salt Lake City, USA), sampled at 30 kHz. Visual stimuli were presented on a Display++ LCD monitor (Cambridge Research Systems, Rochester, UK) with a resolution of 1920 × 1080 px, running at 120 Hz fps.

### Experiment 2: Surgical and experimental procedures

Two male marmosets (cj21 and cj22) were fitted with a titanium head-post to stabilize their head during behavioral training and subsequent recording sessions. The head-post surgery was performed under sterile conditions. Animals were first premedicated with injections of atropine (0.2 mg/kg, i.m.) and diazepam (2 mg/kg, i.m.) 30 minutes prior to the induction of anesthesia with alfaxalone (Alfaxan, 10 mg/kg, i.m., Jurox, Rutherford, Australia). They were then intubated and anesthesia was maintained throughout surgery by inhalation of isoflurane (0.5–3%) in O_2_. Animals were placed in a stereotaxic frame and stabilized using ear bars and a bite bar. The scalp was reflected and the mastoid muscles separated to expose the cranium. Six titanium bone screws (length: 4 mm; diameter: 1.5 mm) were inserted 1–1.5 mm into the skull. The exposed surface of the skull was coated with dental varnish (Copalite; Temrex Corporation), and a thin layer of dental adhesive (Supabond; Parkell) was then applied to the skull to form the perimeter of an acrylic head-cap. The titanium head-post was then positioned on the midline, 3–5mm forward of bregma. Transparent dental acrylic (Ortho-Jet; Lang Dental Mfg. Co.) was then used to encapsulate the base of the head-post and the heads of the bone screws, forming the body of the acrylic head-cap, securing the head-post to the skull. Once the implant was formed, the wound margin was cleaned, and the skin glued to the base of the implant using surgical adhesive (VetBond; 3M). After recovery, animals were acclimated to being head-fixed, and were trained to maintain fixation on a small target. The marmosets underwent a second surgery to chronically implant an electrode array (64 channels, N-Form arrays; Modular Bionics Inc.) in the right V1. The array implantation surgery was performed under sterile conditions with premedication and anesthesia regimes identical to those for head-post surgery.

Extracellular neural signals were amplified, high-pass filtered (0.1 Hz), and sampled at 30 kHz using an Open Ephys data acquisition system (Siegle et al., 2017; OpenEphys, Cambridge, USA). During the recording sessions, animals were seated in a custom-made primate chair with the head-post secured to the chair (Mitchell et al., 2014). Visual stimuli were presented on a VIEWPixx3D LCD monitor (VPixx Technologies Inc.) with a resolution of 1920×1080 px (W x H), running at 100 fps. The stimulus monitor was positioned 48 cm in front of the animal, spanning 57°x32° (W×H) of the visual field. Eye position was tracked monocularly at a rate of 1000 Hz using an Eyelink II video eye tracker (SR Research Inc.). Each monkey completed three sessions of 100 trials on three consecutive days. On every trial, the stationary stimulus patch (see *Visual Stimulation*) appeared for 20 secs. The monkey was required to keep its eye-position within a circular area of 10 deg from the screen center, otherwise the stimulus disappeared and the trial was ended. Liquid reward was given at the end of successfully completed trials (0.02 ml, New Era pump systems, USA). The stimulation trials were randomly interleaved with shorter (3 secs) baseline periods (10 for every session of 100 trials), during which a small fixation target was presented at the center of the screen. On these trials, the monkey was required to maintain its eye position within a circular area of 2.5 deg from the fixation target.

### Visual Stimulation

Visual stimuli were generated in Matlab (The Mathworks, Inc.) and presented using Neurostim *(https://github.com/klabhub/neurostim)* and the Psychophysics toolbox (Brainard, 1997). The main stimulation procedure in both experiments followed the design established in VanRullen and MacDonald (2012), with some adaptations to monkey neurophysiology. The stimulus in both experiments was a circular patch of uniform luminance (diameter, exp. 1: between 3 and 4 deg, adjusted online to cover the receptive field (RF) of the population, exp. 2: always 3 deg) presented on a black background. The patch’s luminance on each trial followed a flattened white noise sequence over time between black and white (exp. 1: 10 s, exp. 2: 20 s duration) with a luminance change on every frame (exp. 1: 120 fps, exp. 2: 100 fps, allowing for stimulus frequencies up to 60 and 50 Hz, respectively). In exp. 1, individual sequences were repeated pseudo-randomly between blocks of 40 or 20 unique trials (monkey cj12: two sessions of 40 sequences x 8 repetitions; monkey cj11: one session à 40 × 8, two sessions à 20 × 8). We found no effect of sequence repetition on any of the measures reported here and thus collapsed data from all recorded trials (N = 640) in both monkeys. In exp. 2, the presented sequences were all independent. The position of the patch in exp. 1 was chosen to best cover the population RF of the recorded units (determined online, adjustments made between recording sessions to account for fixational drift), and was always in the lower right quadrant of the screen. In exp. 2, the patch was centered at [−1.4 −0.2] deg (from screen center, negative values to the bottom & to the left) for both animals. This position was chosen such that the stimulus patch was inside the population RFs approximately at central fixation.

### Analysis of eye-position data (exp. 2)

Eye-position data were analyzed offline. Blinks were detected from the raw eye-tracker data as periods where the pupil area was zero. Subsequently, saccades were detected by applying a threshold criterion (30 deg/sec) to the eye-position velocity traces. Fixations were marked as the time periods between 20 ms after saccade offset up to 20 ms before next saccade onset if eye-position was spatially stable (< 1 deg deviation) in this period. For all analyses in which LFP data was mapped to 2D eye-position (patch-in-RF region estimation, traveling wave fits) we used the instantaneous eye-position, excluding blinks and saccadic periods, resampled to the time-vector of the LFP.

### LFP signal processing

Raw neural data were recorded at 30k samples/sec. To isolate the LFP, the broadband signal was low-pass filtered using an FIR filter with a cutoff-frequency of 200 Hz. For all analyses involving correlation with the stimulus, the signal was low-pass filtered below half the stimulus refresh-rate (anesthetized recordings: 120 Hz, awake: 100 Hz) and then resampled to the time vector of frame presentation times on each trial. Instantaneous phase information was extracted from both signals (stimulus and LFP) by applying a continuous wavelet transform (using the analytic morse-wavelet with γ = 3) to the preprocessed data.

### Analysis of spiking activity

Spiking clusters were initially classified from the continuous data using the unsupervised KiloSort 2 algorithm (Pachitariu et al., 2016). The identified clusters were manually curated and flagged as either noise, multi-unit or single-unit neural signals using the Phy software (developed by Cyrille Rossant; https://github.com/cortex-lab/phy/). Our analysis did not rely on a strict isolation of single units. Therefore, we included all neural clusters, regardless of whether they were classified as multi-or single unit. Data from exp. 1 were sorted by concatenating the two recording sessions for each animal. For exp. 2, we performed the same sorting procedure on the data from each recording day. However, this yielded a very low number of clusters (N < 10), therefore we did not include spiking activity in our analysis of exp. 2.

### Steady-state phase-locking analysis (LFP)

Our main analysis targeted the neural representation of continuous temporal information contained in the stimulus. To quantify this representation without factoring in absolute amplitudes, we reduced both signals (LFP and stimulus sequence) to their phase information and computed an index of the phase-coupling between them (*Steady State Phase Locking* - SSPL), as follows:

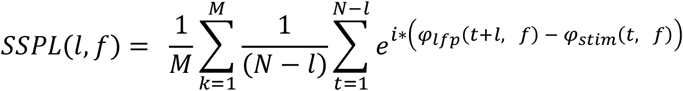

where *φ*_*x*_(*t,f*) is the instantaneous phase of signal *x* at time *t* and frequency *f*. For each lag *l* between LFP and stimulus, the phase differences between the two signals are averaged over time (*t*), and trials (*k*), as vectors of unit length. The resulting SSPL spectrum contains complex values with magnitudes in the range [0 1], with zero and one indicating random and perfect coupling to the stimulus, respectively. The phase angle of each bin represents the average phase difference at that lag and frequency. The inverse transform of the complex SSPL spectrum returns a time-domain impulse response function that is comparable to, e.g., a simple cross-correlation between the two signals, but importantly does not reflect signal amplitudes and their correlation.

For statistical evaluations, we compared SSPL magnitude or phase for single electrode contacts to a bootstrapped random distribution. To this end, we performed the same computation as described above by randomly shuffling the stimulus data in time and drawing with replacement from the resulting set of data. Statistical significance was determined by comparing the observed data to the 95% confidence interval of the random distribution.

To investigate the time-course of LFP-stimulus coupling we derived an index of instantaneous coupling from the global SSPL spectrum for each individual contact:

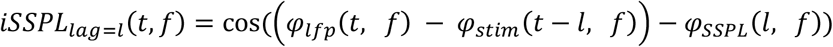

where *φ*_*SSPL*_(*l,f*) is the phase angle of the average SSPL spectrum at lag *l* and frequency *f*. The iSSPL is computed for a single, fixed lag between the signals. The resulting real-valued spectrum (signal time x frequency) represents the degree to which phase differences at that lag are aligned with the average phase difference (i.e., the SSPL phase) for each time x frequency bin.

### Analysis of spike-triggered average stimulus phase (exp. 1)

To quantify the coupling of individual spiking clusters to the stimulus, we relied on the same measure of phase-consistency as for the SSPL analysis in the LFP. Specifically, for a given cluster, we computed the angular mean of the spike-triggered stimulus phase, as follows:

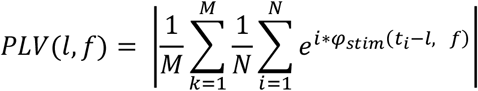

Here, the phase-locking value *PLV(l,f)* represents the consistency with which the spikes (*i* = 1 … *N*) of the cluster followed a certain stimulus phase. It is computed, at each frequency bin *f*, from the angular sum of stimulus phases *φ*_*stim*_ preceding each spike (occurring at time *t*_*i*_) by lag *l*, averaged over trials (*k*).

### Spatial analysis and traveling wave models

In addition to the temporal SSPL analysis we also investigated whether stimulus-coupled information in the LFP was propagated across “remote” cortical regions, i.e., those not covering the instantaneous retinotopic representation of the patch. Based on findings from human EEG studies (Lozano-Soldevilla and VanRullen, 2019; Schwenk et al., 2020) we hypothesized that such a propagation would show in the form of a traveling wave from the retinotopic representation outwards. To test this hypothesis, we extended our analysis of the SSPL response to the spatial domain. Here, we used retinal space, i.e. the relative position of the stimulus to the fovea, as an approximation for the (retinotopic) cortical space in V1. Moreover, since the position of the stimulus patch on the screen was fixed, we could use the instantaneous eye position as an equivalent measure of the patch’s retinal position (the two varying only by the fixed offset of the patch to the screen center). Thus, for each contact, we first mapped the phase differences between the LFP and the stimulus sequence (at each frequency bin) to the instantaneous eye position at stimulus presentation time (in bins of 0.5 deg in both dimensions). From these maps (in screen coordinates), we first estimated the region of eye positions for which the stimulus patch was inside the RF of the local population (*patch-in-RF region*). This estimate was set as a circular region of the same size as the stimulus patch, centered on the position of the peak SSPL response (based on a spatially smoothed map of phase differences).

To isolate any consistent propagation across cortical / retinal space, we then referenced the (unsmoothed) maps of phase differences to the assumed wave center inside the patch-in-RF region (which was therefore set at θ = 0 rad). The analysis region of interest was limited to all positions with sufficient amount of data (min. total fixation time 20 sec/deg^2^), excluding the patch-in-RF region. Additionally, since we had no hypothesis about how the wave would travel between hemispheres, we excluded positions to the ipsilateral (left) side of the patch center. We then fitted a radiating wave model of the re-referenced phase differences to the resulting maps, as follows:

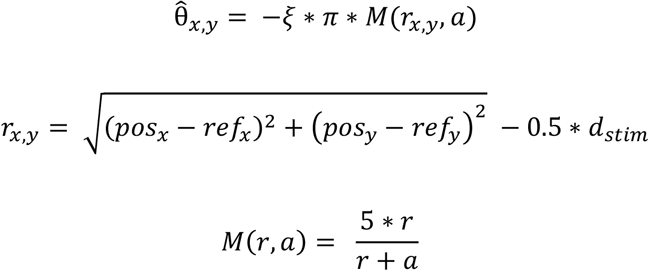

where 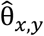 is the predicted phase difference for eye position *pos*_*x,y*_, and *ref*_*x,y*_ is the eye position where patch center and the estimated RF are aligned (patch-in-RF region center). Phase differences are modeled here as a function of the radial distance *r* of the eye position to the outer edge of the stimulus patch (patch center + half the patch diameter *d*_*stim*_). Since foveal and parafoveal retinal regions are magnified in cortical space in primate V1, the instantaneous frequency of a traveling wave (assuming a constant frequency across the cortex) would be inversely proportional to eccentricity when mapped onto the retinal space. To account for this, the distance *r* is converted to approximate cortical distance through the magnification factor *M*(*r*). The parameters for the cortical magnification functions differ between species, and, in the marmoset, are best described by an exponential term multiplied by the scaled inverse of the eccentricity, allowing 3 free parameters (Chaplin et al., 2013). Given the low spatial resolution in our data we chose only an approximate model for *M*, with one free parameter (*a*) and the scaling factor (*k = 5*) approximated based on Chaplin et al. (2013).

The second free parameter of our model was the spatial frequency of the wave *ξ* (proportional to its propagation speed). The best fit was determined from an iterative search within the 2D space of all possible *ξ* ∈ [0.1 2] (in units of radial distance in mm^−1^ * cycles) and *a* ∈ [0.3 0.9], by maximizing the cosine-weighted sum of the circular residuals between actual and predicted values of θ, defined as:

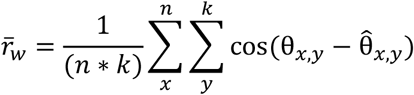

As a standardized measure to compare model fits across contacts and monkeys, we used the square of the circular correlation coefficient (ρ_*cc*_) (following the analysis in Zhang et al. 2018):

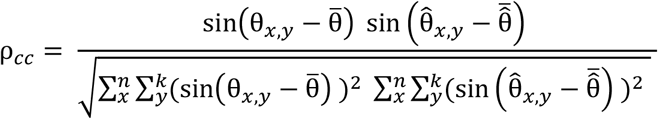

where, as before, θ is the observed and 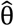 the predicted phase difference, and averaging 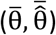 is performed across all positions in *x,y*. Statistical significance of model fits for individual contacts was determined by comparing the fit to a distribution of 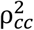 values resulting from fits to bootstrapped maps (based as before on a phase-time-shifted stimulus sequence).

In a separate step, we also tested for each fit if it allowed for a better prediction of the phase difference maps than reached by a mean-phase approximation (null model, no phase gradient). For this comparison, we used the weighted sum of residuals 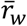 defined above.

## Results

We recorded neural responses to broadband visual flicker in V1 of marmoset monkeys under general anesthesia (Exp. 1) and in the awake, behaving state (Exp. 2).

### Exp. 1: Anesthetized state

With the recordings from anesthetized animals, our goal was to characterize the passive response properties of V1 populations to random luminance sequences. In brief, two animals (cj11 and cj12) were stimulated with broadband visual flicker sequences of 10s duration, while neural data was collected from a 96-channel Utah array in V1. The luminance stimulus (a circular patch) was positioned inside the population RF.

### Local field potentials

We first analyzed stimulus responses in the LFP. In the trial average, all contacts showed a clear response to the onset of the stimulus sequence (Fig. 1A). Peak latencies (to the first, smaller peak) of this response were in the expected range of V1 latencies (Yu et al., 2012) (Fig. 1B), with a slight difference between animals (median latency, *cj11*: 62.5 ms; *cj12*: 56.2 ms). As our main interest was on the continuous response to the stimulus, we limited our further analysis to the steady state after this initial onset response, i.e., 250 ms post sequence onset to trial end. From this time period, we extracted phase information from the stimulus sequence and the recorded LFP on each trial and computed a time-frequency-resolved index of the coupling between both signals (*Steady State Phase Locking -* SSPL). The results of this analysis are shown in Fig. 1C, for a single contact in each of the two monkeys. The SSPL response is defined as a complex spectrum across lag (time) and frequency, of which the magnitude (top panels in Fig. 1C) represents the strength of coupling to the stimulus. In both monkeys, the LFP phase exhibited broadband responses to stimulus frequencies between 3 and 30-40 Hz. Coupling was stable for a single oscillatory cycle only, i.e., did not show selective entrainment at single frequency bands. This is also evident when observing the inverse transform of the SSPL spectrum (bottom panel in Fig. 1C for the same contacts as above, representing the corresponding time-domain impulse responses), which generally showed a similar shape as the responses to sequence onset (Fig. 1A). Indeed, the peak latency of the onset response predicted the phase of the SSPL response reliably across contacts (Fig. 1D, circular-linear correlation significant at p<0.001 (corrected) in both monkeys). Fig. 1E shows a summary of the frequency distributions of coupling strength across contacts. Notably, the average response in monkey cj12 showed a global peak in the theta range (7.2 Hz, dashed vertical line) that was not visible in the other monkey.

**Figure 1.**
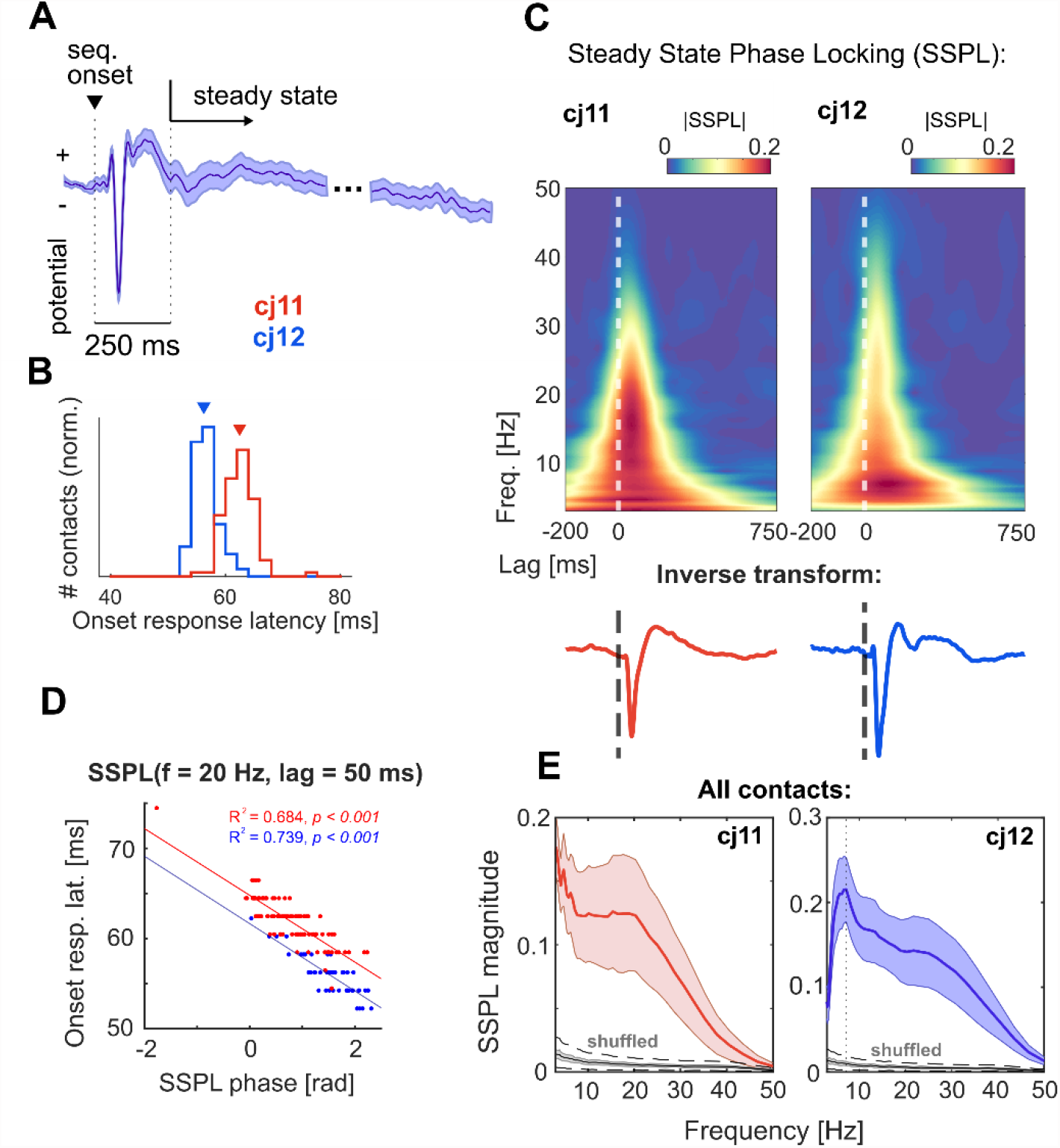
Steady-state phase locking (SSPL) responses in the LFP of anesthetized marmosets. **A**: Evoked response to sequence onset. The trace shown is the trial-average from a single contact in monkey cj12 (SEM in the shaded area). For the subsequent analyses, the LFP in each trial was cut to a time-window starting 250 ms after sequence onset (*steady-state*). **B**: Distribution (across contacts) of peak latencies for the response to sequence onset. Latencies relate to the first positive deflection in the trial-average evoked response (cf. A). **C:** top panels, SSPL responses computed across trials for a single contact in each monkey. The color axes indicate the magnitude of the complex SSPL spectrum, representing strength of coupling between LFP and stimulus at a given lag and frequency. Bottom panels show the corresponding time-domain representations of the same responses (arbitrarily scaled). **D**: Correlation between SSPL phase and onset response latencies across contacts. Colors indicate data from the different monkeys as before. Phase values were evaluated at 50 ms lag and a frequency of 20 Hz. **E:** Average frequency spectrum of the SSPL response magnitude across all V1 contacts. Colored curves show mean +/-1SD in the shaded areas. Grey curves show the SSPL obtained for a random shuffling of the stimulus data (see Methods), with the shaded area representing SD across trials, and the dashed black lines showing the average 95% CI of a single contact.

### Spiking activity

In addition to the continuous LFP response, we were also interested in how the stimulus information was represented in V1 local spiking activity. To this end, we isolated spiking clusters (single-or multi-unit, N = 56 in cj11 and N = 53 in cj12) and calculated their coupling to the stimulus spectrum analogously to the SSPL response in the LFP. Fig. 2A shows an example of a single cluster’s response to sequence onset, characteristic of the population response in both monkeys. Spiking activity returned to pre-stimulus baseline levels (or even below) within a few hundred ms after a brief transient burst following stimulus onset (log ratio of steady-state to baseline firing rate, pooled: t(108) = −1.678, p = 0.096). Within this steady state, we quantified each cluster’s coupling to the stimulus by calculating the spike-triggered average stimulus phase at varying lag and frequency. Fig. 2B shows the resulting response spectra for four exemplary clusters. Each cluster showed a unique frequency distribution in its coupling to stimulus phase, with peaks ranging between the theta and the lower gamma band. Notably, the distribution was bimodal in some clusters (e.g., second and fourth panel in Fig. 2B). This may either reflect the presence of multiple cells within the cluster with different properties, or complex response behavior within a single cell. The variety of responses between clusters is again also evident from the time-domain impulse response (bottom panels in Fig 2B, all magnified to the first 200 ms). Here, each response curve (colored) is overlaid with the same cluster’s response to sequence onset (PSTH, grey). This comparison revealed a consistent shift towards lower latencies from the onset response to the steady-state (Fig. 2C), which was statistically significant in both monkeys (cj11: mean shift, M = −4.726 ms, t(32) = 11.917, p < 0.001; cj12: M = - 9.158, t(39) = 2.486, p = 0.035; p-values corrected for multiple comparisons).

**Figure 2.**
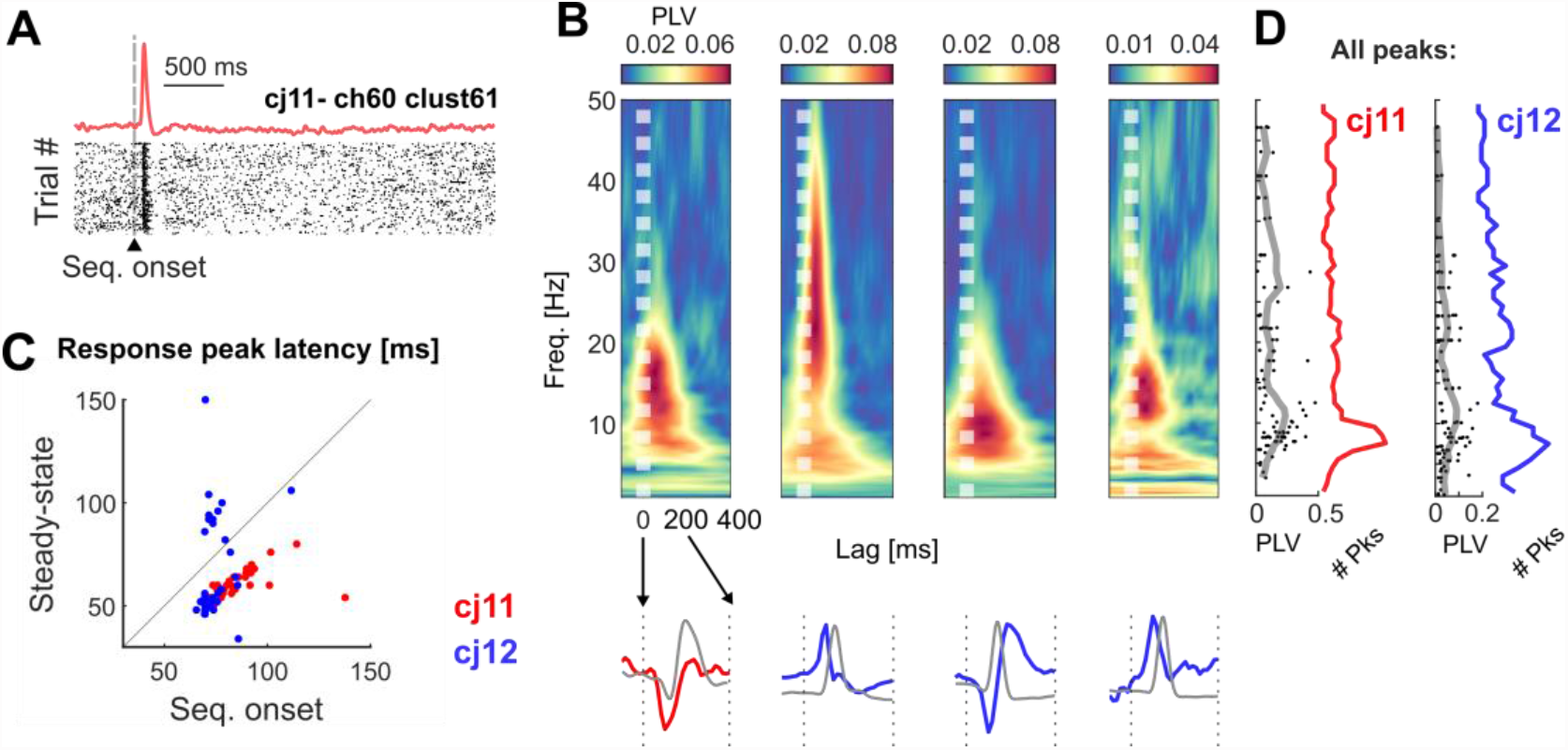
Steady-state coupling of V1 spiking clusters to the stimulus sequence (anesthetized). **A:** Sequence onset response in a single cluster. The raster plot shows spikes in single trials, with the corresponding PSTH plotted above. Note that the time axis does not include the full 10s sequence presented on each trial. **B:** Steady-state spike-stimulus coupling in four selected clusters. The top panels show the spike-triggered average stimulus-phase spectra, with strength of phase coupling indicated by the color axis (PLV). The time axis refers to the lag from stimulus to the triggering spike (i.e. flipped compared to a classical spike-triggered average). The bottom panels show the corresponding inverse transforms (colored traces), overlaid with the sequence onset response for the same cluster (grey). Both responses are enhanced to show only the first 200 ms lag. **C:** Comparison of peak latencies for onset and steady-state responses. Latencies were significantly reduced in the steady-state in both monkeys. Only clusters for which the peaks in both responses had the same sign were included in this analysis. **D:** Population distribution of peak coupling frequencies. Single dots translate to frequency peaks in the responses that were significantly greater than chance level (more than one per cluster possible). Colored curves show the histograms, grey curves the moving average of the peak PLV. Red and blue colors refer to data from the two different monkeys throughout the figure.

To approximate the summed response of the population as a whole, we extracted all peaks with significant coupling strength (as compared to a bootstrapped random distribution) from each cluster’s response, and then pooled across clusters (Fig. 2D). The resulting distribution showed a higher density in the upper theta range, peaking at the same frequency bin (7.5 Hz) in both monkeys. However, on average, coupling strength (grey curves, moving average PLV) was highest for peaks at slightly higher frequencies, in the alpha range. The variability in peak coupling frequency between clusters was not explained by differences in firing rate (evaluated for the largest PLV-peak for each spiking cluster), neither from the pre-stimulus baseline nor the steady state (all correlations between firing rate and frequency of peak PLV, R^2^ < 0.06, *n*.*s*.). This suggests that the preferred coupling frequencies of each cluster in the steady-state constitute a unique response property of the underlying neuron(s), similar to a classical temporal-frequency tuning curve.

### Exp. 2: Awake, behaving state

After establishing the passive response properties of V1 populations to broadband flicker, we next asked how the stimulus representation would be modulated under natural conditions, i.e., during active vision. In the second experiment, we used the same basic stimulation procedure as before (central circular patch, modulated in a random luminance sequence on each trial, here for 20 s duration) while recording from laminar probes in area V1 of two awake marmosets (cj21 and cj22). The monkeys were allowed to view the stimulus display freely (within a circular region 20 deg in diameter centered on the screen), but on average eye-position remained just to the right of the stimulus patch (cj21, weighted center [horz./vert.] [1.11 −0.01] deg) or diagonally above it (cj22, [1.92 2.85] deg) (Fig. 3A, left panels). Fixation durations showed the expected left-skewed distribution (Fig. 3A, right), with median durations at 302 ms (cj21) and 197.5 ms (cj22). To analyze the main stimulus response, we calculated the SSPL spectrum from the LFP in the same way as for Exp. 1, first collapsed over the full trial. Fig. 3B shows the result of this for single contacts in each monkey. Similar to Exp. 1 (Fig. 1C), the responses were limited to a single or a few cycles, but here the spectra showed a marked separation into two distinct frequency bands. We denote these as the low- and high-frequency component (LF/HF) in the following. The bimodality in the frequency distribution was highly consistent across contacts (Fig. 3E), and the peak frequencies of the two components were also within a narrow range unique to each monkey (peak of the average, cj21: 9.473 Hz (LF) and 23.326 Hz (HF); cj22: 13.397 Hz (LF) and 23.326 Hz (HF)). In contrast to this, the power of the raw LFP signal (i.e., non-phase locked, Fig. 3F) showed a prominent peak in the beta range (peaking at the same frequency bin in both monkeys, f_max_ = 13.4 Hz). This distribution was consistent across stimulation (solid curves) and baseline periods (blank screen, dashed curves) (aside from differences in amplitude, which we did not consider further because fixation behavior was not comparable between the two conditions).

**Figure 3.**
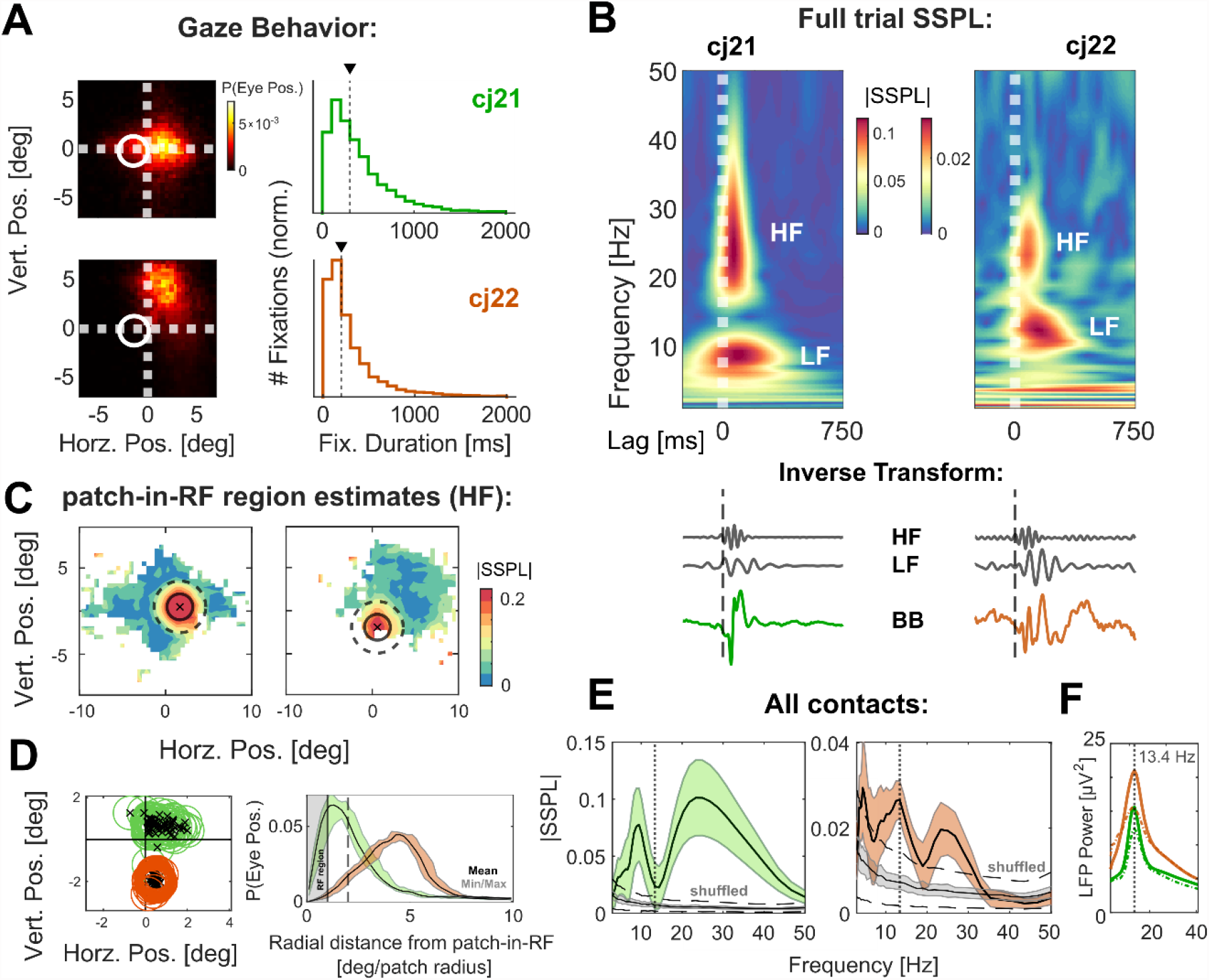
Steady-state LFP-stimulus coupling in V1 of awake marmosets. Green and orange colors refer to data from the two monkeys (cj21 and cj22, respectively) throughout the figure. **A:** Summary of eye-movement behavior during the steady state (all trials collapsed) in each monkey. Left panels: spatial distribution of eye positions (in screen coordinates). The location of the stimulus patch is shown as a white circle. Right: histograms of fixation durations. Dashed vertical lines mark median durations (cj21: 302 ms; cj22: 197.5 ms). **B:** top panels: SSPL responses for a single contact in each monkey; analogous to Fig. 1C. Responses were computed from the full trial (excluding only the first 250 ms), i.e. not accounting for eye-position. Bottom: time-domain representations of the same responses, again analogous to Fig. 1C (BB: broadband), but here plotted also for the band-limited transforms of LF and HF components. **C:** Maps of SSPL magnitude as a function of eye position (HF component, single contacts). These maps were used to estimate the patch-in-RF region (eye positions for which the patch was covering the RF) of each contact. Patch-in-RF region center and area (1 patch radius) are marked by cross and solid circle, the dashed circle represents an area of 2 patch radii. **D:** Left: summary of patch-in-RF region estimates for all 64 contacts in each monkey. Right: distribution of eye-positions expressed in radial distance from the patch-in-RF region center. Shaded area represent minimum and maximum across contacts. Solid and dashed vertical lines represent 1 and 2 patch radii as in C. **E:** Mean frequency spectra of SSPL magnitude across contacts, using the same conventions as in Fig. 1E. Dashed vertical lines correspond to the global peak in LFP power (13.4 Hz in both monkeys). **F:** Global power-spectrum of the raw LFP (non-phase-locked, mean across contacts) during stimulation (solid curves) and without (blank screen).

Since eye position was not fixed, the RFs of the local population at each contact were not aligned with the stimulus patch throughout the trial. We estimated the responses’ spatial profile by re-computing the HF component at each contact as a function of retinal space (using eye position at stimulus presentation time as an equivalent measure). As expected, the resulting maps (Fig. 3C; in screen-coordinates; single contacts for each monkey) revealed strongest coupling within an area of the same size as the patch (solid circle). Note that, to yield sufficient mean vector lengths, the maps in Fig. 3C were smoothed in the complex domain using a box of 1 patch radius width (1.5 deg), resulting in the visible halo (dashed circle, 2 patch radii). From this, we estimated a *patch-in-RF* region for each contact, defined as the range of eye-positions for which the patch was inside the assumed RF of the local population (Fig. 3D, left panel; also in screen coordinates). These estimates allowed us to distinguish for each local population between direct (retinotopic) and indirect (remote) stimulation. The alignment of fixation preference and patch-in-RF regions (Fig. 3D) confirmed the opposite pattern of data availability between monkeys, explaining the large differences in SSPL magnitude in the trial-average. As a consequence, we limited our analysis of retinotopic processing dynamics to monkey cj21.

### Temporal dynamics of the SSPL response

Here, we were specifically interested in the dynamics of the SSPL response over the time course of a single fixation. To this end, we derived an index of the instantaneous coupling from the average response for each contact (iSSPL, see Methods for details), and compared this measure across fixations. The average time-frequency spectrum for all fixations > 200 ms (Fig. 4A, single contact) clearly demonstrates that coupling evolved as a gradient with a linear increase beginning shortly after fixation onset. For stimulus information presented within a brief perisaccadic time window, coupling at higher frequencies was notably suppressed (note that the response delay is fixed here to 70 ms). However, the overall frequency distribution (LF and HF split) did not change throughout the fixation. We therefore quantified coupling dynamics again for LF and HF separately and used the component average time-courses to compare fixations depending on the direction of the preceding saccade, i.e., whether it brought the stimulus into the population RF or kept it there (Fig. 4B).

**Fig. 4:**
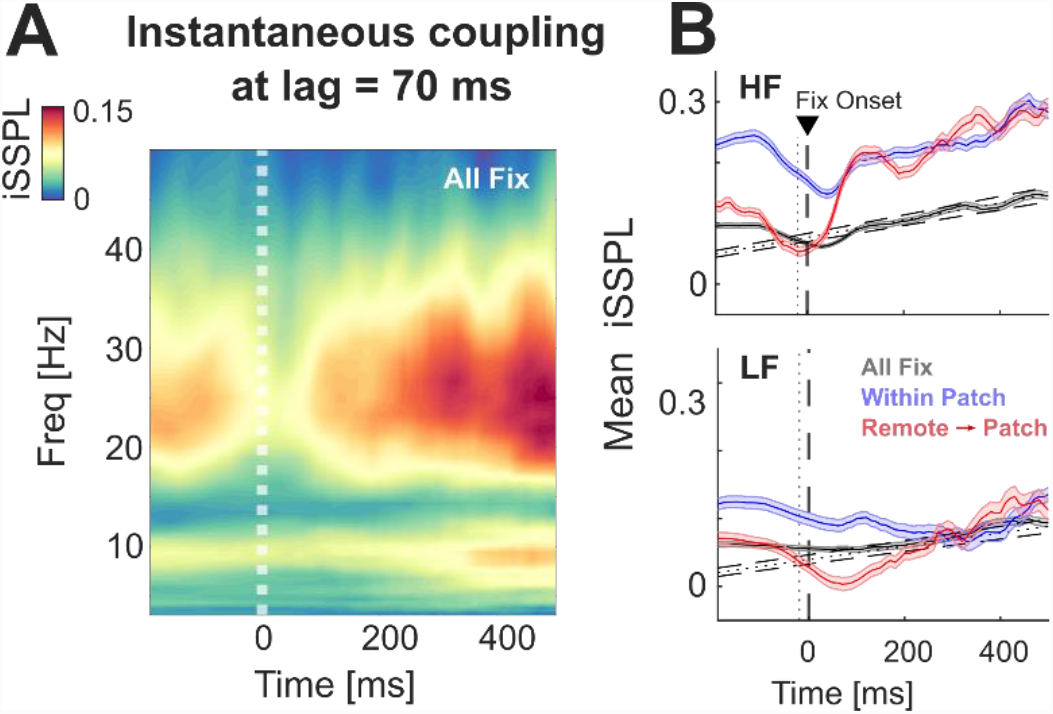
SSPL dynamics across the saccade-fixation cycle. Data from a single monkey (cj21), including fixations of > 200 ms duration. **A:** Instantaneous coupling across the average time-course of a saccade and following fixation (for a single contact). Time axis is aligned to fixation onset (t = 0 ms) and coupling was evaluated relating to stimulus information presented at t-70 ms. **B:** Time-courses of mean iSSPL within the HF and LF components, for the same alignment as in A (average across contacts). Data from patch-in-RF fixations were split between preceding saccades bringing the patch newly into the RF (red) and those maintaining patch-RF alignment (blue). Dashed lines show a linear fit to the time-course of all fixations (mean +/-average 95% CI across contacts).

This analysis revealed that coupling dynamics were similar following saccades bringing the patch into the RF (red curves) and those where it was inside before and after (blue). Interestingly, however, in the latter case coupling was still above baseline-levels around the saccade, indicating that processing was continuous at least to a certain degree. We confirmed the time-dependent increase in coupling strength by fitting linear slopes to the time-courses for single contacts (dashed line), which showed significant linear trends in 60 (93.75%) and 59 (92.19%) of the 64 contacts for LF and HF, respectively (compared to bootstrapped random distributions; mean slopes in ΔiSSPL*sec^-1^: r_LF_ = 0.099, r_HF_ = 0.142). We also verified that the observed increase was not a result of a systematic phase-shift in the coupling between LFP and stimulus.

### Responses outside the RF: traveling wave models

Our final analysis was directed at possible activity modulation outside the RF. For the above analyses of stimulus representation inside the RF, the SSPL response could be described by a constant phase difference within the boundaries defined by the patch. For non-RF locations on the other hand, SSPL phase should vary across retinal space since the responses would likely reach those regions by lateral propagation. Specifically, based on previous studies (Lozano-Soldevilla and VanRullen, 2019; Schwenk et al., 2020) we hypothesized that this variation would show in the form of radially traveling waves centered on the retinotopic patch representation, selectively for the lower frequencies. To test this, we fitted the observed phase differences (i.e., phase angles of the maps in Fig. 3C) with a simple radiating wave model. The spatial frequency of the wave was set as a free parameter and the fitted region limited to eye positions outside our estimated patch-in-RF regions, for each contact’s LF and HF component separately. We evaluated the fitted models in two steps. First, statistical significance of each fit was determined by comparing it to a random distribution. Fig. 5A shows the summary of this analysis, where the ratio of explained circular variance (ρ^2^_cc_) is plotted against peak (temporal) frequency. In both monkeys, fits reached higher ρ^2^_cc_ values than expected by chance on a subset of contacts (p < 0.05, uncorrected; cj21, LF: 8 (12.5%), HF: 18 (28.13%); cj22, LF: 31 (48.44 %), HF: 22 (34.38 %)).

**Fig. 5:**
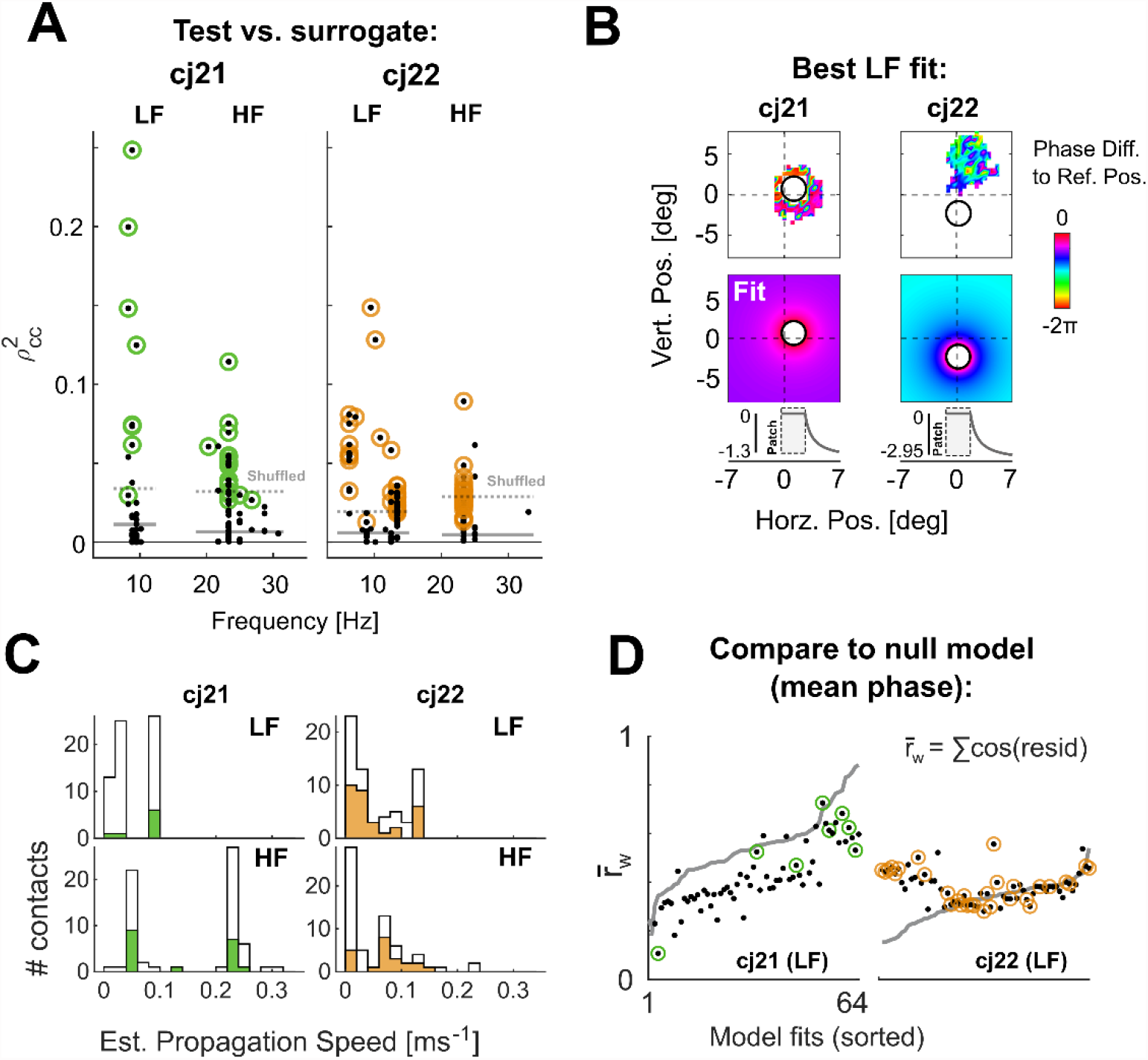
Traveling wave fits of the LFP-stimulus phase differences outside the patch-in-RF region. **A:** Summary of best model fits for each contact and frequency component. Each dot represents the fit for a single contact, plotted as the ratio of explained variance, ρ^2^_cc_, against frequency. Colored circles mark ρ^2^_cc_ values greater than the 95% CI of a bootstrapped random distribution (mean and average CI across contacts indicated by horizontal solid and dashed lines, respectively). **B:** Phase difference maps (top panels: observed, bottom: predicted) for the contacts with the best LF fit in each monkey. Differences are re-referenced to zero at the assumed wave center. The grey curves show a radial cross-section of the predicted phase-difference (patch-in-RF region in the shaded area). Note that position/distance in all panels is plotted in retinal coordinates, while phases are modeled as a linear function of assumed cortical distance. **C:** Estimated propagation speeds, derived from the combination of spatial (ξ) and temporal frequency of each model. Histograms show all 64 contacts in the white, and significant fits at p<0.05 (uncorr.) in the colored areas. **D:** Comparison of the traveling wave fit (LF only) to the mean-phase null model for each contact. Scatterplots show the cosine-weighted sum of circular residuals between actual and predicted phases (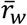,higher values indicate a better fit) of the traveling wave model, with the grey curve representing 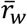 of the null model (sorted in ascending order). Colored circles mark significance as in A.

Since our expectation was on the lower frequencies, we scrutinized the obtained LF models further (examples in Fig. 5B, showing the highest ρ^2^_cc_ across contacts). Notably, a considerable amount of wave fits predicted very low propagation speeds (< 0.05 ms^-1^, Fig. 5C). This result may indicate that the model type was inappropriate, specifically if the actual phase differences are better described by a constant offset between inside and outside the patch-in-RF region. Therefore, in a second step, we tested the validity of the model type by comparing each fit to a null model of a constant phase difference (Fig. 5D). Here, we found that most model fits for cj22, and all for cj21, performed at about the same level of or worse than the null model (grey curves in Fig. 5D). For the remaining fits in cj22 (N = 9, i.e. those statistically significant and valid over the null model), propagation speeds were all < 0.01 ms^-1^. While it is possible that these contacts indeed showed a consistent propagation of stimulus phase, we see it as more likely that they represent either random effects (carried by the multiple comparisons across contacts) or other inhomogeneities in the underlying maps (e.g., bad estimates of the RF-region). In summary, we did not find clear evidence that stimulus information was propagated across non-RF regions in the form of (temporally consistent and spatially regular) traveling waves in the LFP.

## Discussion

Our results provide the first systematic description of how broadband visual flicker is represented in primate V1. In the anesthetized animal, we found broadband coupling to the stimulus phase in the LFP, and more selective coupling in individual spiking clusters. Our LFP recordings in awake animals showed selective coupling in distinct frequency bands unique to each monkey. This response was confined to the retinotopic representation of the stimulus in V1, and showed a systematic increase over the time-course of single fixations.

### Temporal frequency selection in the steady state

Data from both experiments show that, within a few hundred ms following stimulus onset, the responses reach a steady state in which the LFP couples its phase to that of the stimulus sequence. Steady-state responses to visual flicker have been described before using single frequencies (e.g. in cat area 17, Rager and Singer 1998). Broadband sequences, on the other hand, have been used extensively with human EEG (VanRullen and MacDonald, 2012; Chang et al., 2017; Gulbinaite et al., 2017; Benedetto et al., 2018; Lozano-Soldevilla and VanRullen, 2019; Schwenk et al., 2020). Our study provides a first step towards linking these results to cellular-level V1 activity. The selective frequency tuning observed in awake animals suggests that this paradigm can capture properties that would not be evident from mapping single frequencies at a time. Of the presented broadband stimulus, only two distinct bands were represented reliably in the LFP (alpha, 8-13 Hz; beta, 20-35 Hz). This frequency selection likely represents a correlate of active vision: responses in the anesthetized animal were broadband, demonstrating that synchronized LFP and spiking activity would in principle allow for coupling to frequencies outside the above components. The possible functional role of this bimodal selection remains unclear. One explanation may be that specific frequency bands are actively disengaged from the stimulus because they are engaged in other processing. In one monkey, the component split corresponded to the global beta peak in the raw LFP power spectrum (cj21, Fig. 3E, F). This peak was similarly present in the other monkey, although here, it overlapped with the LF component. Beta-band oscillations in the visual cortex have been hypothesized to reflect top-down signaling of task- and behavioral contexts to modulate bottom-up processing (Engel and Fries, 2010; Richter et al., 2018). In our experiments, without a behavioral task and long trial durations (20 secs), this signal may reflect behavioral idleness and/or prediction about the continued presence of the stimulus. It is unclear if and how this (stimulus-unrelated) beta oscillation would relate to the drop in stimulus coupling. It is possible that distinct local populations (stimulus-coupled and top-down beta) contribute concurrently to the LFP, whereas high phase-stability of the latter amplifies their contribution artificially. However, as noted before, this antagonistic relationship is supported by the data from only one monkey.

Another explanation for the observed tuning is a split-by-design. That is, each component may represent information that is extracted from the stimulus for a different purpose. The LF component could represent a similar narrow-band filter as the perceptual echo response in the human EEG, which has been associated with active rhythmic sampling (VanRullen, 2016; Benedetto et al., 2018) and temporal prediction of upcoming visual input (Chang et al., 2017; Alamia and VanRullen, 2019). These theories could be tested at the neural level in future studies by employing the methods used here. For instance, the rhythmic sampling account would predict that spiking activity (either locally or at remote cortical locations) be modulated by the phase of the evoked lower-frequency oscillation, e.g. by altering response gain (Haegens et al., 2011; Haegens and Zion Golumbic, 2018; Davis et al., 2020) or temporal integration windows (Chota and VanRullen, 2019; Chota et al., 2021). Our results show that spiking activity may be analyzed using the same analysis tools as for the LFP, yielding complementary results. Temporal frequency tuning is a well-known property of V1 neurons, which show individual preferences for single-frequency stimuli (macaque: Foster et al., 1985; Hawken et al., 1996; marmoset: Yu et al., 2010). We interpret our results from spiking clusters in the anesthetized animal as reflecting a similar property. Clusters showed consistent preferences for frequency bands, covering the full frequency range across the population. Notably, this tuning emerged in a sparse steady state with firing rates at or below baseline-levels, indicating an adapted spike-time coding regime (Prescott and Sejnowski 2008).

### Temporal coupling dynamics

In the awake response of one monkey we found that LFP coupling strength increased with fixation duration. This result likely reflects two different mechanisms. Saccades bringing the stimulus into the RF are correlated with a transient increase in firing rate within the first few hundred milliseconds following fixation onset, suppressing stable phase-coupling of local neurons to the stimulus. In addition, saccades that re-position the stimulus inside the RF may evoke a similar suppression, since fixation onsets are followed by brief transients of high excitability and a reset of LFP phases in V1, facilitating new visual input (Rajkai et al. 2008).

Both mechanisms highlight that the visual system may essentially alternate between two modes: one optimized to process larger changes to the visual scenery, the other to evaluate continuous temporal information. The animals’ gaze behavior suggests a clear bias towards the former mode, in line with ecological demands (Mitchell et al., 2014; Mitchell and Leopold, 2015). Interestingly, human observers making saccadic decisions based on the perceived brightness of a similar random luminance flicker relied disproportionately on the first 100 ms of the 1 s sequence (Ludwig et al., 2005), indicating that sensory weights may be similarly biased away from the steady state. Given the short fixation durations in our data, it remains unclear at which duration the steady state is fully reached (i.e., when coupling strength plateaus). Human EEG studies employing the same stimulus used longer fixations (3.125 secs in Chang et al., 2017; 6.25 secs in VanRullen and MacDonald, 2012). It may be interesting to explore the dynamics of the temporal response function across the saccade-fixation cycle also in the human EEG, in particular for the perceptual echoes (as discussed below).

### Spatial response profiles

We found that, in retinal space, LFP stimulus-coupling was tightly confined to the retinotopic representation of the stimulus patch. This is in line with standing estimates of the cortical volume that contributes to the LFP signal (approx. 200 μm in the absence of correlation; Katzner et al. 2009; Lindén et al. 2011). Our analysis of the surrounding positions did not provide evidence that information was propagated laterally across the cortex, as hypothesized from the human perceptual echoes (VanRullen and MacDonald, 2012).

We see several possible explanations for this result. First, as discussed above, the echo response may rely on longer fixation durations to emerge. The observed LF responses were within the extended alpha range. If this component corresponds to a resonance frequency of V1 neurons (Herrmann, 2001), surrounding cortical regions could be selectively brought to oscillate in relative phase with the RF at this frequency. This mechanism could produce traveling waves if the resonance frequency itself shows a gradient across the cortex (theory of weakly coupled oscillators; Ermentrout and Kleinfeld 2001; Zhang et al. 2018). However, unlike direct lateral propagation this would require a prolonged steady state to allow local oscillators to couple their phases. To illustrate, the median fixation duration in cj21 was approx. 300 ms, which, accounting for response delay, allows for only two alpha cycles to be completed before the next saccade. Additionally, surrounding areas may be less susceptible to pick up the evoked oscillation within the first few hundred ms of a fixation because their network-state is transiently synchronized (Lefebvre et al., 2017) as a result of post-saccadic phase-resets (Rajkai et al., 2008).

An alternative explanation for the absence of lateral propagation could be the small size of the marmoset cortex. Previous studies have shown similar speeds across species for lateral propagation of evoked responses and spontaneous traveling waves, of up to ∼0.8 ms^-1^ (Bringuier et al., 1999; Muller et al., 2014; Zhang et al., 2018; Davis et al., 2020). At the same speed, a single cycle covers the same retinal distance in the marmoset faster than in macaques or humans. For higher speeds, the resulting phase gradient may not have been distinguishable from the mean-phase null model in our data.

Related to this, the use of a laminar array (implanted vertically in the cortex) presents a limitation of the present study. Our analysis had to rely on a reconstruction of cortical space through retinal space (recorded *sequentially*); thus, the phase maps used to evaluate the traveling wave hypothesis were not based on *simultaneously* recorded signals. Instantaneous phase gradients over cortical space would allow for a more sensitive wave detection, and also the detection of waves that occur irregularly over time or with variable spatial frequency (Townsend et al., 2015; Townsend and Gong, 2018). However, measures of stimulus-coupling that are based on phase-differences, as is the case for the SSPL, typically rely heavily on averaging over time (i.e. fixations and/or trials). Dedicated investigations of the cortical propagation of stimulus-coupled activity will thus need to develop measures indicating how irregular wave events detected from single trial activity relate systematically to the stimulus.

Taken together, our results are insufficient to rule out cortical traveling waves in the steady-state response. However, they show that, if these waves are present, they are not detectable as stationary phase-gradients in retinal space for short fixation durations.

## Acknowledgements

This work was supported by Deutsche Forschungsgemeinschaft (RU 1847; IRTG-1901), EU (PLATYPUS) (all to FB), the Australian Research Council (DE180100344 to MH; DP200100179 to NP and MH; DP210103865 to MR and SC; DP210101042 to MR and EZ) and by the National Health and Medical Research Council of Australia (APP1194206 to MR, APP1185442 to MH and SC, APP1120667 to NP). We also thank Janssen-Cilag for the donation of sufentanil citrate. We are grateful to Rufin VanRullen for helpful comments on the results.

